# Deciphering the multi-scale mechanisms of *Tephrosia purpurea* against polycystic ovarian syndrome (PCOS) and its major psychiatric comorbidities: studies from network-pharmacological perspective

**DOI:** 10.1101/785048

**Authors:** Neha Choudhary, Shilpa Choudhary, Vikram Singh

## Abstract

*Tephrosia purpurea (T. purpurea)*, a plant belonging to Fabaceae (pea) family, is a well-known Ayurvedic herb and is commonly known as Sarapunkha in traditional Indian medicinal system. Described as “*Sarwa wranvishapaka*”, *i.e.* having capability to heal all types of wounds, it is particularly recognized for its usage in splenomegaly. Towards exploring the comprehensive effects of *T. purpurea* against polycystic ovarian syndrome (PCOS), its phytochemicals were extensively reviewed and their network pharmacology evaluation is carried out in this study. The complex regulatory potential of its 76 phytochemicals (PCs) against PCOS are enquired by developing and analyzing high confidence tripartite networks of protein targets of each phytochemical at both pathway and disease association scales. We also developed a high-confidence human PPI sub-network specific to PCOS, explored its modular architecture and probed the drug-like phytochemicals (DPCs) having multi-module regulatory potential. The proteins belonging to endocrine system were identified as major targets of the PCs. The study reports 30 DPCs based on ADMET and drug-like properties. Multi-targeting and synergistic capacities of the 12 DPCs against 10 protein targets were identified and evaluated using molecular docking and interaction analyses. The study concludes by highlighting a couple of DPCs as potential sources of PCOS regulators.

## Introduction

Polycystic ovarian syndrome (PCOS), a genetic condition of ovarian dysfunction is the most common form of endocrinopathy with global prevalence of 6-10% amongst women of reproductive age (Goodarzi, Dumesic, Chazenbalk, & Azziz, 2011). The complex and heterogeneous nature of PCOS is associated with both non-reproductive and reproductive morbidity. Apart from anovulation, amenorrhea, hirsutism (hyperandrogenism) and infertility observed in the women with PCOS, increased BMI is prevalent in 30-70% of cases (Vrbikova & Hainer, 2009). Long term morbidity factors associated with PCOS include sub-fertility, obstetrical complications, diabetes mellitus, cardiovascular disease, malignancy and psychiatry in some cases.

Stein and Leventhal, in 1935, were the first to describe PCOS as a combination of signs and symptoms related with the androgen excess and dysfunction of ovaries, hence called as stein-levithel syndrome (Endocrinology and 2018). In 2012, National Institutes of Health (NIH) sponsored “Evidence-Based Methodology Workshop on Polycystic Ovary Syndrome” recommended a new name to this condition *i.e.* PCOS (R. et al., 2016). Both genetic and environmental factors play a major role in disease development. The complex nature of PCOS and its associated comorbidities undermine the management of this disease during the treatment procedure. The comorbid relationship of PCOS is with a variety of mental and metabolic diseases that includes symptoms of anxiety, depression, bipolar disorders, diabetes, cardiovascular complications *etc*.

Since the treatment of PCOS is mainly focused on meliorating the symptoms associated with hyperandrogenism, variety of adverse-effect profiles have been encountered by the prevailing pharmacological therapies (Domecq et al., 2013). It is being recognized that therapies focusing on ameliorating the comorbid relationship should also be given key importance while designing new therapeutic strategies. Thus, a search of therapeutically effective procedures with reduced side effects and the multi-targeting property is of urgent need and an utmost priority for the researchers working in this area. In recent years, systems biology based drug-design approach of “Network-pharmacology” has gathered significant attention for unrevealing the multi-targeting framework of various small molecules for their therapeutic relevance in a very high-throughput manner. Various traditional herbs like *Piper longum* (Choudhary & Singh, 2018) and formulations like triphala (Chandran, Mehendale, Tillu, & Patwardhan, 2015) *etc*. have been explored using this approach to provide a mechanistic understanding to the traditional knowledge and providing novel ways to derive therapeutically active compounds for the drug-discovery. Also, augmentation of the concepts of network-pharmacology with CADD (computer-aided drug-design) are proposed to be of great help towards delineating the therapeutic action of natural compounds against various disease and disorders of mankind (Choudhary & Singh, 2019).

Ayurveda, referred to as “Science of Life” is the traditional Indian system of medicine which aims to promote and sustain health with a balanced lifestyle. Effective in dealing with the disease at its prevention, diagnosis and treatment stage, an extensive knowledgebase of Ayurveda holds a detailed description of various medicinal plants and their associated formulations for the disease management. *Tephrosia purpurea,* a plant belonging to Fabaceac (pea) family is a well-known plant of traditional Indian Ayurvedic literature, commonly referred to as Sarapunkha. In Ayurveda, this plant is described as “*Sarwa wranvishapaka*” meaning it possesses the ability to heal all types of wounds and is highly acclaimed for its beneficial effects towards maintaining female reproductive health (Goswami, Dash, & Dash, 2011; Thakor & Patel, 2014). In Ayurevdic texts, various parts of this plant and its associated formulations are described for the treatment of diverse range of diseases and disorders (Sharma, 2005).

*T. purpurea* is found as a common wasteland weed and grows in poor soils. The plant is perennial herbaceous in nature with highly branched and sube3rect stem structure (Sharma et al., 2013). The dried herb is effectively used as a tonic laxative, de-obstruent and also in complications associated with asthma and urinary disorder (Bhardwaj, Shrivastava, & Shrivastava, 2017). Medical properties of this plant include antiallergic activity, anti-lipid peroxidative, hepatoprotective activity, immunomodulatory, antimicrobial as well as chemopreventive activity. The plant is also well recognized for the management of complications linked to liver cirrhosis and splenomegaly (Bhardwaj et al., 2017) and is reported to be effective against diseases of spleen, heart, blood, kidney and liver (Ajitha et al., 2014).

In this study, we have first reviewed the phytochemicals present in *T. purpurea* by an extensive literature mining and search in public databases. The collected phytochemicals are clustered on the basis of their chemical classification and subjected for ADMET property evaluation to screen putative drug-like candidates (*i.e.* DPCs). The drugability of the compounds were further assessed by performing their structural similarity with the existing drugs of DrugBank. The polypharmacological action of phytochemicals present in *T. purpurea* was inferred using the approach of network pharmacology where polypharmacological activity of a compound was estimated on the basis of number of their protein targets. The functional relevance of the protein targets have been identified using human pathway mapping, disease-association and module participation. High confidence DPC-protein target pairs associated with PCOS were predicted and subjected for the molecular docking and interaction analysis. The series of steps followed in this study are represented in **Figure 1**.

**Fig. 1:**
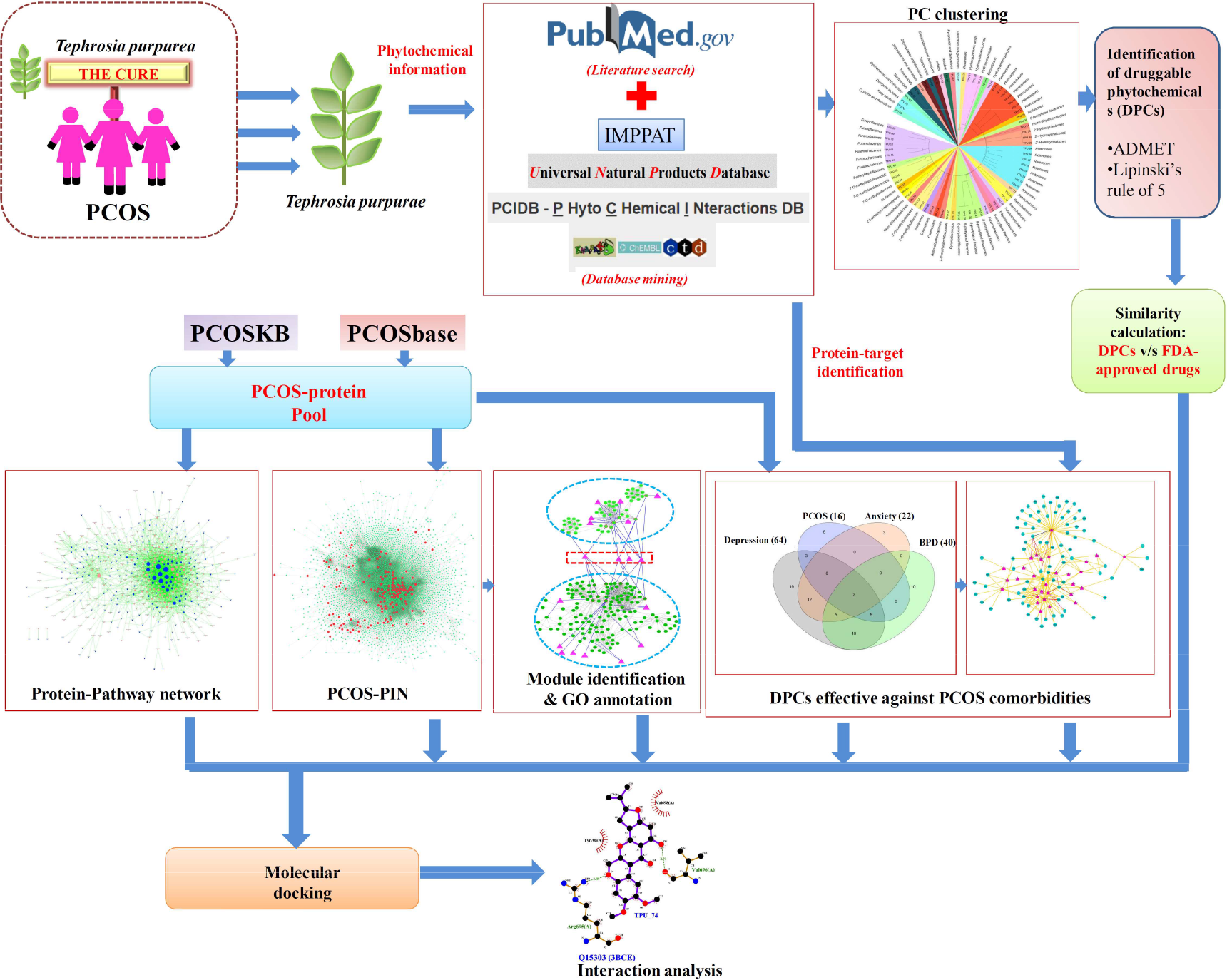
The detailed workflow of present study on network pharmacological evaluation of *T. purpurea*.

## 2. Materials and methods

### 2.1 Data information and collection

A list of phytochemicals reported in *T. purpurea* was compiled using database sources available in public domain and manually curating the PubMed articles. Four public databases, namely, (i) Dr. Duke’s phytochemical database (Duke, 2017), (ii) PCIDB (Phytochemical interaction database) (http://www.genome.jp/db/pcidb), (iii) TCMSP (Traditional Chinese Medicine Systems Pharmacology) (Ru et al., 2014), and (iv) IMPPAT (Indian Medicinal Plants, And Therapeutics) (Mohanraj et al., 2018) were searched for retrieving the information of phytochemicals present in *T. purpurea.*

Also, articles in PubMed, an archive of biomedical and life science research manuscripts, were manually inspected to collect the relevant publications for enlisting the phytochemicals of *T. purpurea.* The phytochemicals compiled from both, literature survey and database mining were examined to remove duplicate entries. Two chemical databases *i.e.* PubChem (Bolton, Wang, Thiessen, & Bryant, 2008) and ChEMBL (Bento et al., 2014) were used to obtain the chemical information of the phytochemicals reported in *T. purpurea.*

### 2.2 Clustering and chemical distribution of phytochemicals

A chemical class was assigned to each phytochemical using ClassyFire, an annotated and rapid chemical classification method which uses the information of chemical structure features for hierarchical classification. (Djoumbou Feunang et al., 2016). Clustering of phytochemicals was performed using clustering toolbox of ChemMine tools. A hierarchical clustering algorithm was opted for searching compound similarities based on Tanimoto coefficient and atom-pair based descriptors.

Based on the Lipinski’s rule of five, a chemical compound bearing molecular weight (MW) < 500 daltons; number of hydrogen bond acceptor (HA_n_) < 10; number of hydrogen bond donor (HD_n_) and logP value < 5 is likely to be a candidate drug molecule (Lipinski, Lombardo, Dominy, & Feeney, 1997). For this, the calculation of the ADMET (Absorption, Distribution, Metabolism, Excretion, Toxicity) properties were performed using pKCSM based on the graph-based signatures (Pires, Blundell, & Ascher, 2015). Only phytochemicals following the Lipinski’s rule were considered for the subsequent analysis and refereed as putative drug-like phytochemicals *i.e.* DPCs. The ADMET values of 30 DPCs are given in the **Supplementary Table-1**.

### 2.3 PCOS-gene library

The proteins associated with PCOS were collected from two databases, namely, 1) PCOSKB, a web-based online knowledgebase containing the manually curated information of 241 PCOS-associated genes, SNPs, associated diseases, pathway and GO association (Joseph, Barai, Bhujbalrao, & Idicula-Thomas, 2015). 2) PCOSbase, having a collection of PCOS-related genes and providing information of 8,185 genes compiled from online sources (which includes OMIM, MalaCards, DisGeNET, DISEASES and DGA) and expression studies (Afiqah-Aleng, Harun, Muhammad, Azlan, & Mohamed-Hussein, 2017). The gene-list obtained from both the sources was compiled and duplicate entries were removed to obtain a unique set of 8,199 genes. This dataset of 8,199 genes constitutes the PCOS-gene library (**Supplementary Table-2**). The collected genes were mapped to their respective Uniprot Ids, and 7,972 (list-1) Uniprot Ids were returned for these 8,199 genes.

### 2.4 Target identification

The potential protein targets of the phytochemicals of *T. purpurea* were obtained from three database sources, namely, **(1)** STITCH 5.0 (Search Tool for the interaction of chemicals and protein), that integrates the information about interactions from metabolic pathways, crystal structures, binding experiments and drug-target relationships (Szklarczyk et al., 2016). The confidence score of ≥ 0.4 was used to screen high confidence protein targets of the query molecule. **(2)** Binding DB, that is a database of experimentally known binding affinities of protein-ligand pairs (Liu, Lin, Wen, Jorissen, & Gilson, 2007). The similarity score of ≥ 0.85 was used to retrieve binding data of the query molecule. **(3)** SwissTargetPrediction, that returns top-15 protein targets based on the combined score of 2D and 3D similarity measures between the query molecule and its most similar ligands (Gfeller et al., 2014).

### 2.5 Similarity index calculation

The chemical similarity between the phytochemicals of *T. purpurea* and the drugs present in DrugBank was assessed on the basis of Tanimoto coefficient (*T*_*c*_) calculated using OpenBabel v2.4.1 (O’Boyle et al., 2011). The structural comparison between the two compounds was performed using FP2; a path based molecular fingerprints which index linear fragments of the molecule from 1-7 atoms where molecular fingerprints help to encode the chemical structure of the compounds in the form of binary digits. *T*_*c*_ between two chemical compounds X and Y is given by,

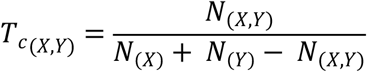

where, *N*_(*X*)_ and *N*_(*Y*)_ are the number of molecular fingerprints in compounds X and Y, respectively and *N*_(*X*,*Y*)_ is the number of molecular fingerprints present in both X and Y (Gohlke et al., 2015; Willett, Barnard, & Downs, 1998). The value of *T*_*c*_ ranges from 0 to 1, with 1 representing maximum and 0 representing no similarity.

For the comparison, PCs passing the drug-likeliness criterion (*i.e.* 30 DPCs) and approved drugs of DrugBank were considered for the analysis. Tanimoto score of DPC-Drug pairs between 30 DPCs and 2,384 approved drugs are given in **Supplementary Table 3**.

### 2.6 Network construction and analysis

STRING v11.0, an integrated database of the protein-protein interaction data was used to identify interaction among PCOS proteins (Szklarczyk et al., 2015). Only the interactions limited to *Homo sapiens* with the confidence interaction score ≥ 0.9 was used to construct the high confidence human PPI-subnetwork specific to PCOS. Molecular COmplex DEtection algorithm; MCODE was used to organize the obtained PPI into functional modules (Bader & Hogue, 2003). Module represents a set of proteins which tend to cluster together and work in a coordinative manner to perform a particular function (Barabási, Gulbahce, & Loscalzo, 2011). The biological relevance of proteins constituting a module was detected using the functional annotation tool provided by DAVID Bioinformatics Resources 6.8 (Dennis et al., 2003).

To examine the poly-pharmacological actions of various phytochemicals, networks involving associations among the phytochemicals (PCs), protein-targets (PTs) of PCs, biochemical-pathways (BPs) and disease-association (DAs) of PTs were constructed and studied using Cytoscape v3.2.1 (Shannon et al., 2003). The knowledge of biological pathways with which the identified protein targets are associated was obtained from KEGG database (Kyoto encyclopedia of genes and genome) (Ogata et al., 1999). A database having comprehensive gene-disease associations called DisGeNET (Piñero et al., 2017) was used to investigate the role of PCs against various disease classes.

### 2.7 *In-silico* molecular docking and interaction studies

AutoDock was used to predict all the favorable conformations of ligand at the active sites of protein targets. This technique combines simulated annealing and a grid-based method for conformation searching (Goodsell, Morris, & Olson, 1996). Docking studies were performed using Lamarckian genetic algorithm. Ligplot+ was used for obtaining the 2D representations of molecular interaction in protein-ligand complexes (Wallace et al., 1995).

## 3. Results and discussion

### 3.1 Phytochemical dataset preparation

A comprehensive list of phytochemicals present in *T. purpurea* was prepared using extensive literature review and mining of public-domain databases having information about natural products. A total of 76 phytochemicals could be found and each of them was allotted with a unique identifier. Detailed information of all the phytochemicals is presented in Table 1.

**Table 1:**
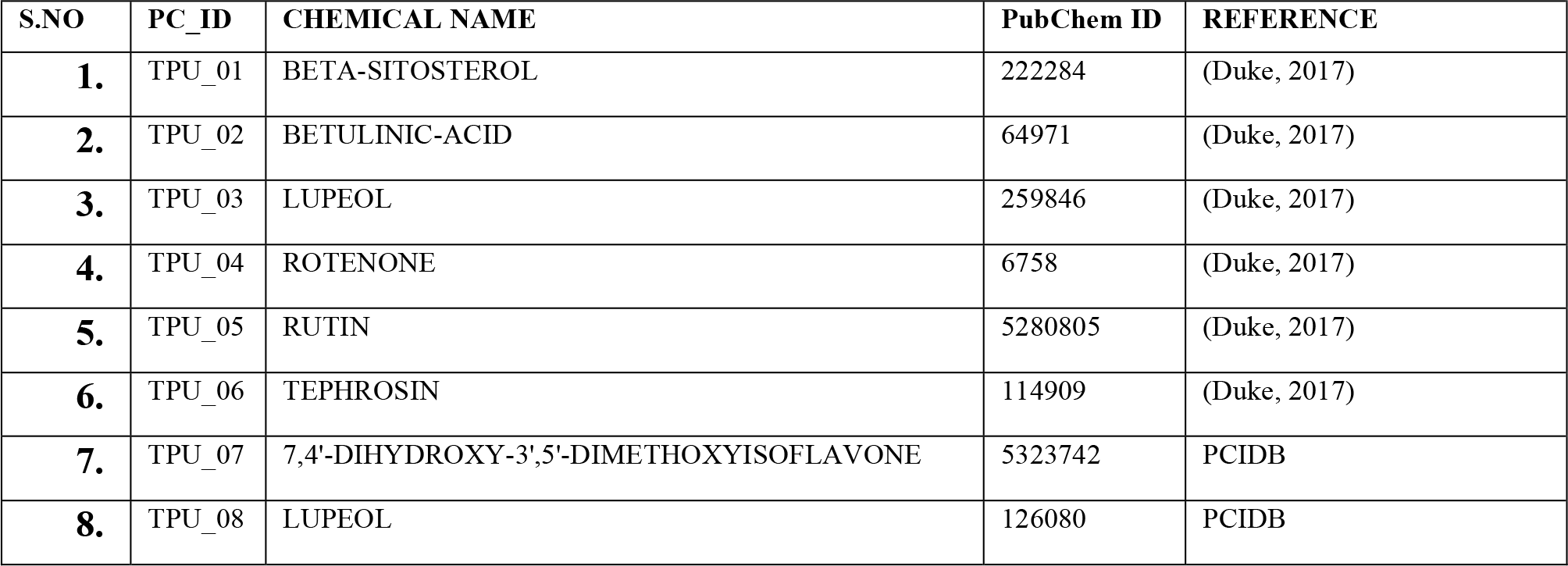
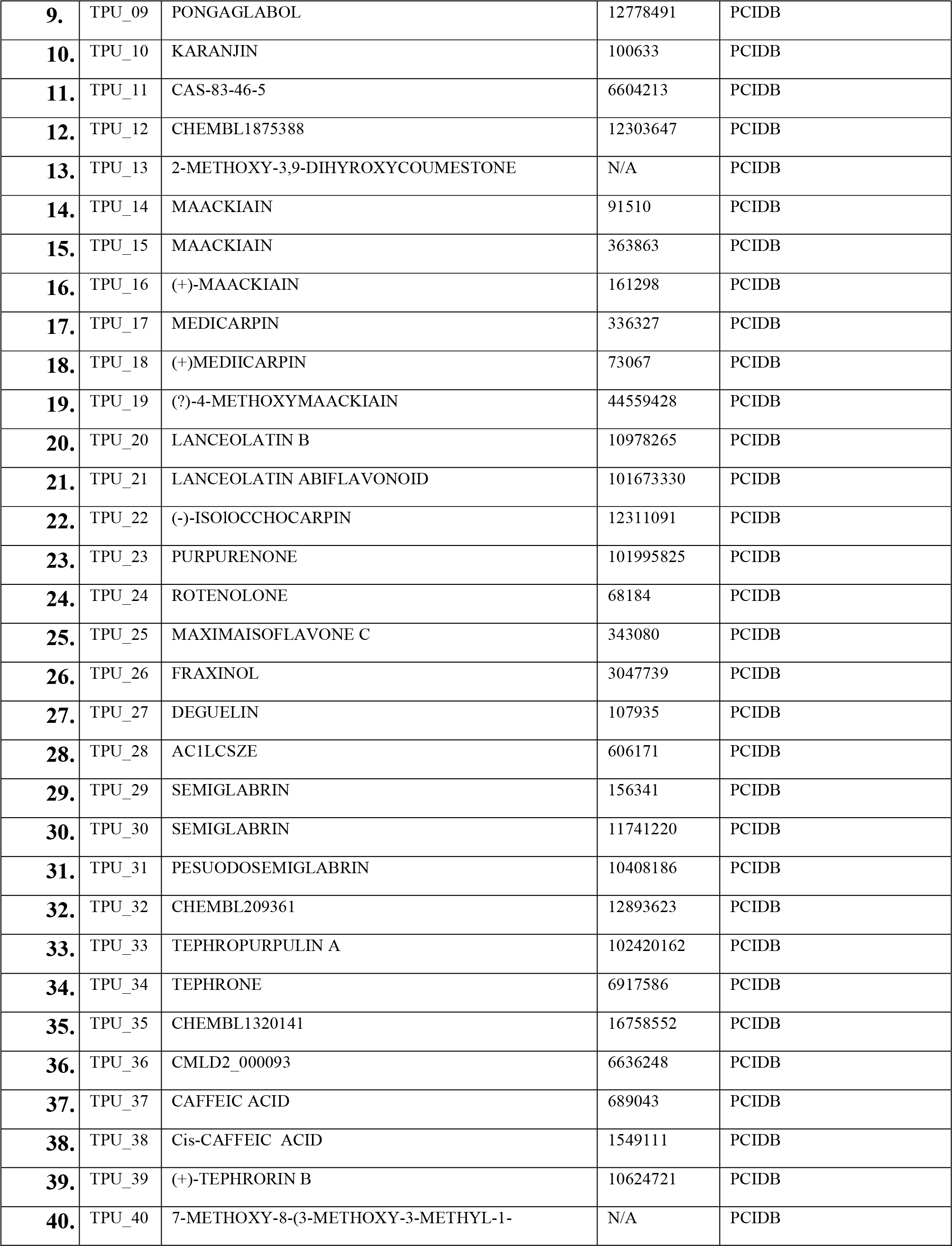
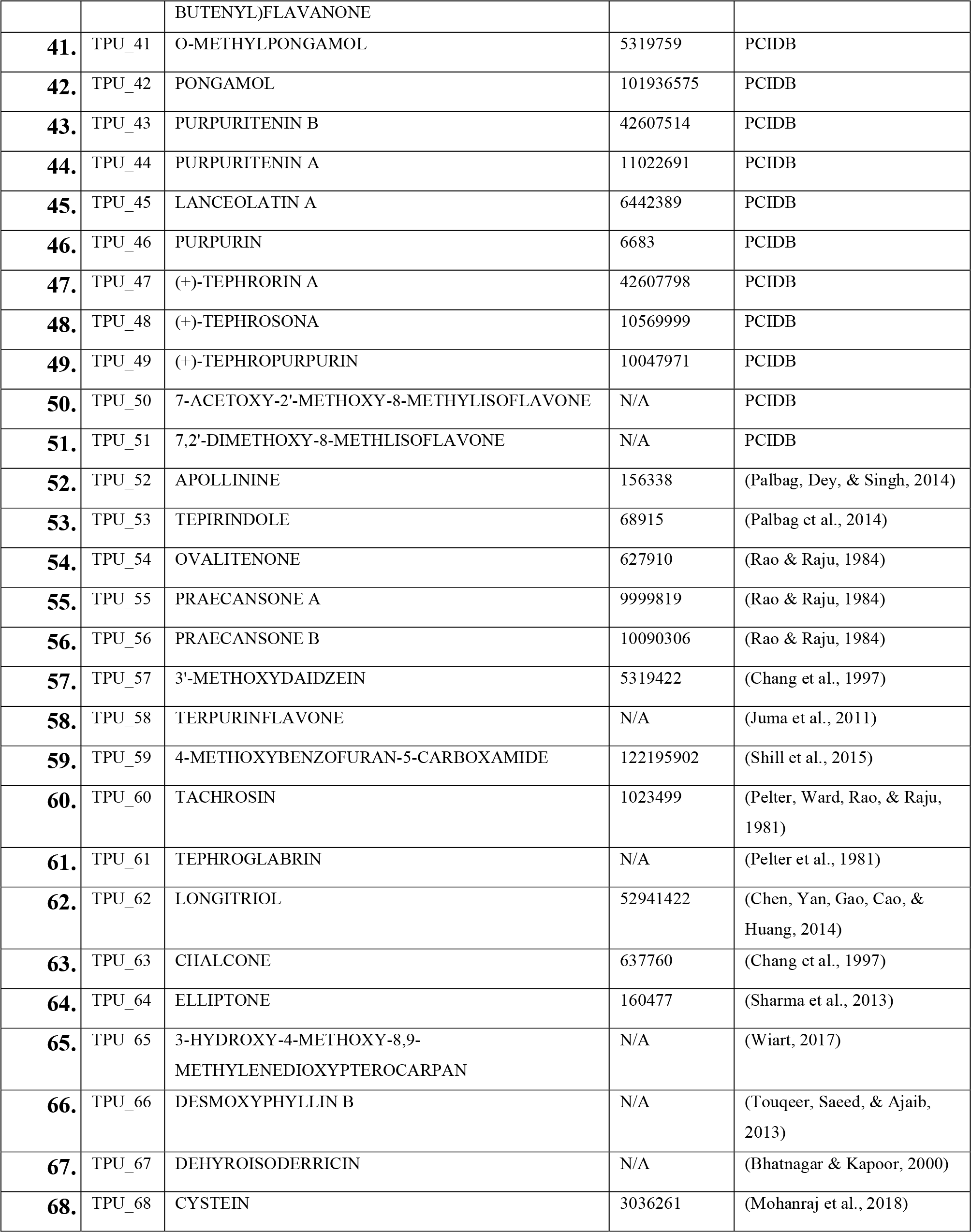
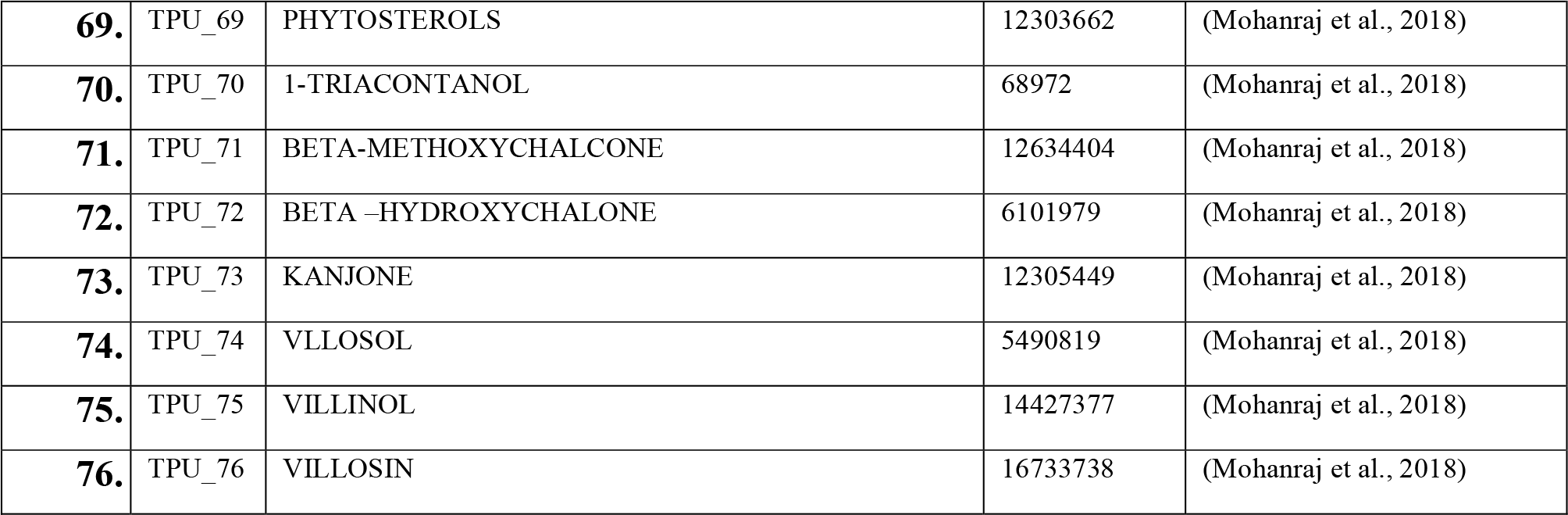
List of the phytochemicals identified in *T. purpurea*

### 3.2 Clustering and chemical distribution of phytochemicals of *T. purpurea*

The PCs are found to be dispersed among 32 chemical classes, out of which the classes of 8-prenylated flavones and pterocarpans are shared by the maximum number of PCs **(Figure 2A)**.

**Fig. 2.**
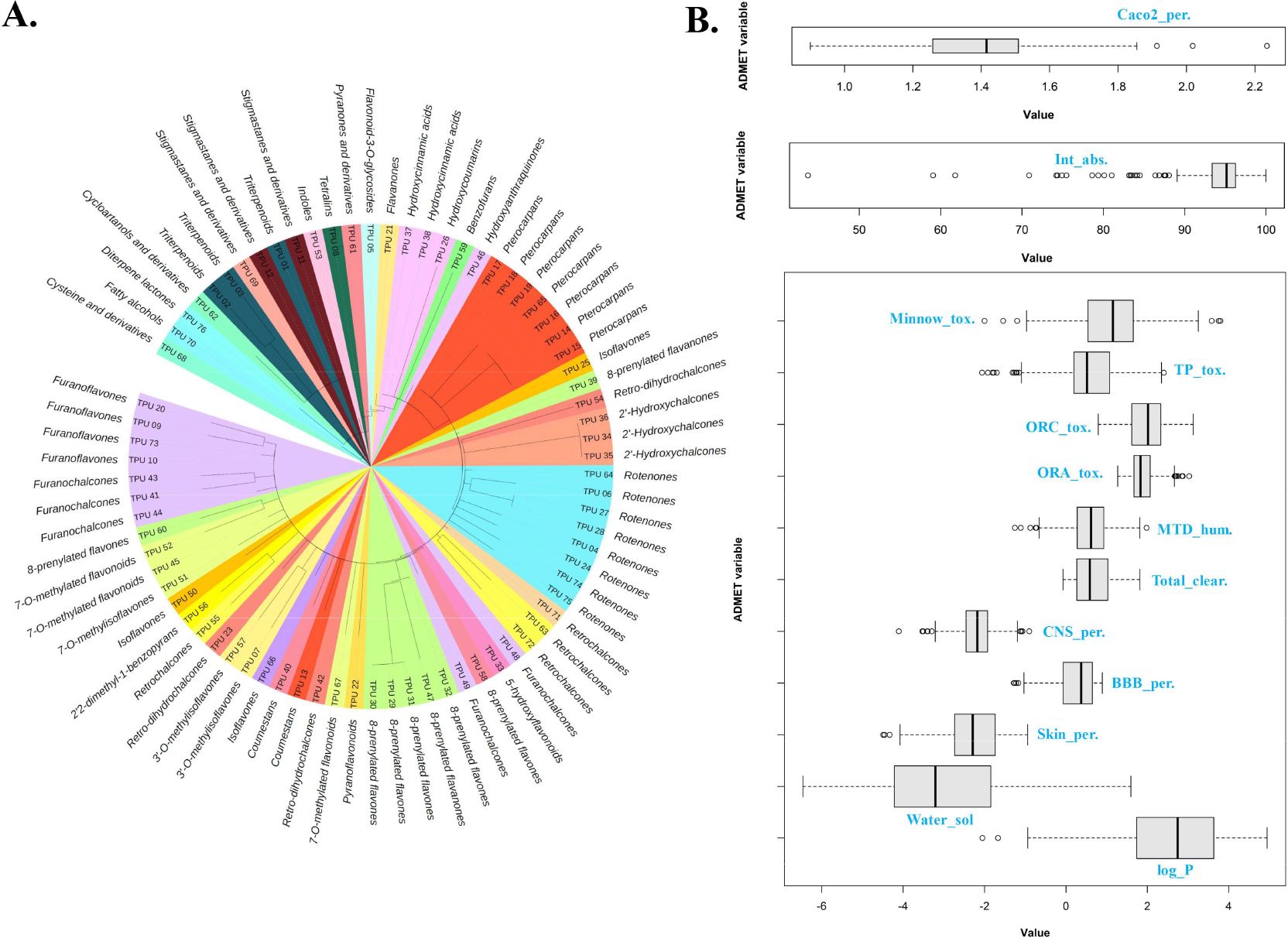
**A: Clustering and distribution of phytochemicals of *T. Purpurea***, The 76 phytochemicals (PCs) of *T. purpurea* are clustered in a hierarchical manner based on the Tanimoto-coefficient and atom-pair descriptors using ChemMine tools. Clustering of the compounds is represented in the form of a tree layout where outer circles represent the chemical class of the PCs. The 76 PCs are found to be classified into 32 chemical classes where each class is represented by a unique colour code. **B: ADMET properties of DPCs of *T. purpurea***, The 13 ADMET properties of 30 DPCs of *T. purpurea* are represented in box and whisker plot. These properties were obtained from pkCSM web server and are as follows; Caco-2 cell permeability (Caco2_per.), intestinal absorption (Int_abs.), minnow toxicity (Minnow_tox), *T. pyriformis* toxicity (TP_tox), oral rat chronic-toxicity (ORC_tox.), oral rat acute-toxicity (ORA_tox.), maximim-tolerated dose in humans (MTD_hum.), total clearance (Total_clear.), central-nervous system permeability (CNS_per.), blood-brain barrier permeability (BBB_per.), skin permeability (Skin_per.), water solubility (Water_sol) and octanal-water partition coefficient (log_P).

Various phytochemical studies provide evidence of the presence of prenylated flavones in *T. purpurea,* especially investigation based on the aerial part (Hegazy, Abd El-Razek, Nagashima, Asakawa, & Paré, 2009). Also, the PCs belonging to similar chemical classes tend to cluster together. The chemical classification of each PC of *T. purpurea* is detailed out in **Supplementary Table-4**.

Many compounds were not able to meet the Lipinski’s rule of five criteria, and only 30 of 76 PCs confirmed the rule. The screened PCs at this level were considered as drug-like phytochemicals (DPCs) and were considered for the further network pharmacological analysis. The ADMET values of 30 DPCs, refereed as DPCs (as mentioned in Section 2.2) are presented in the form of the box and whisker plot **(Figure 2B)**. The list of 30 DPCs and their ADMET values can be checked in the **Supplementary Table-1**.

### 3.3 Phytochemical-drugs similarity network (PC-DR network)

In order to select active compounds in the *T. purpurea* plant based on the PCs similarity with approved drugs of DrugBank, Tanimoto coefficients were calculated. The similarity between molecules is represented in the form of a network, where an edge is drawn between a PC and a drug only if the similarity between them was more than 40% (*i.e.* TC ≥ 0.4). For that purpose, PCs earlier passing the drug-likeliness criterion (*i.e.* 30 DPCs) were considered at this step. Out of the 30 DPCs, 27 are found to have the 40% similarity condition with the 75 drugs. The chemical similarity between these 27 DPCs and 75 drugs is represented in the form of similarity network (PC-DR), with the network size of 102 nodes and 211 edges (**Figure 3; Supplementary Table-5**). Detailed examination of the similarity network helped us to identify that the TPU_04 shares a similarity with 30 out of 75 drugs in the PC-DR network. Also, the maximum similarity of 0.6 exists between DB00553 and TPU_26. TPU_04; Rotenone, classified as moderately hazardous class II pesticide by the WHO is known for its broad-spectrum insecticidal and herbicidal properties (Organization, 2010). This naturally occurring compound has gained massive attention from the scientific community as an investigational drug towards cancer treatment due to its mitochondrial dysfunction ability (Heinz et al., 2017).

**Fig. 3:**
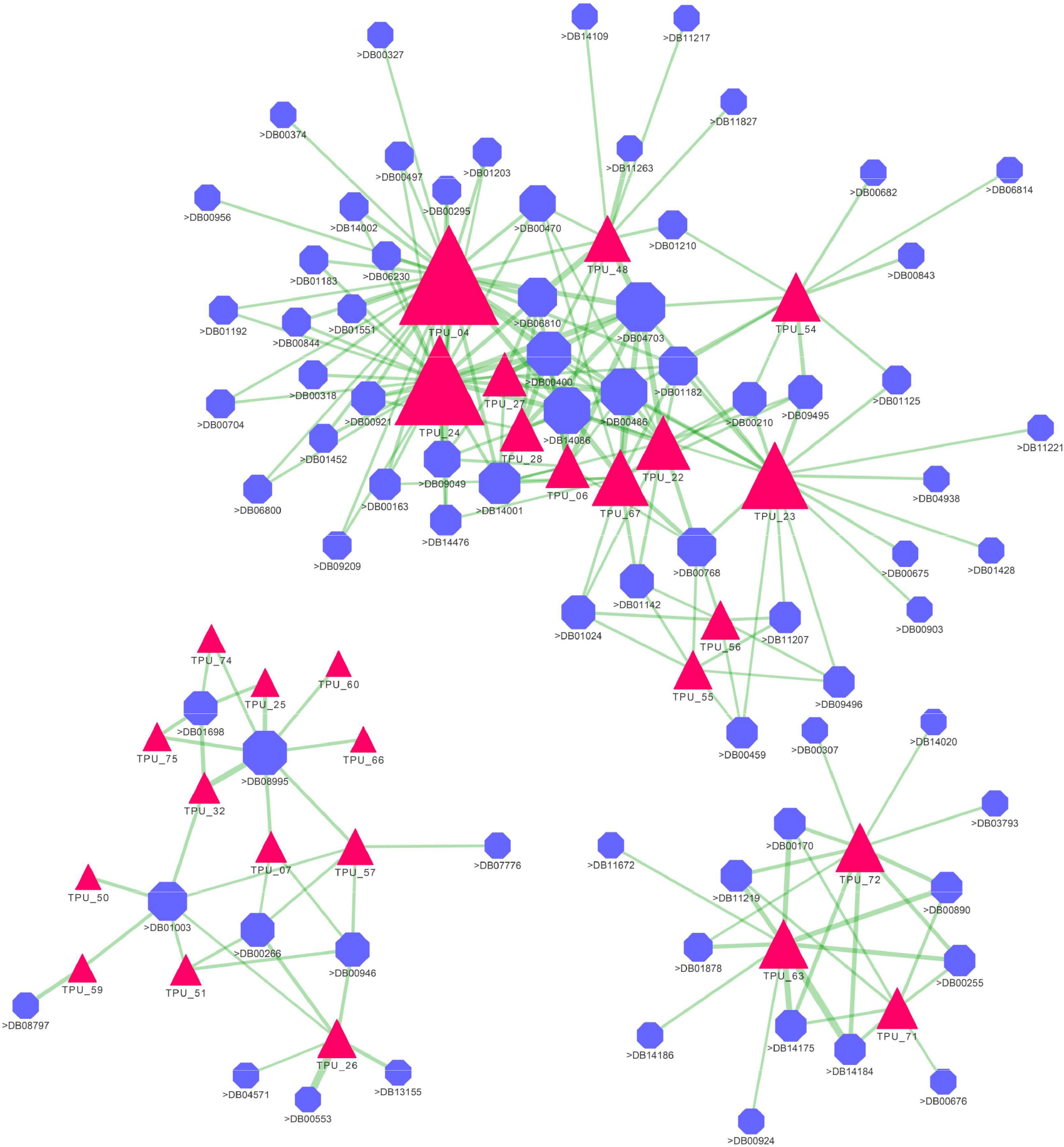
Drug-like phytochemicals – DrugBank compounds (DPC-DBC) similarity network, The DPC-DBC network represents the Tanimoto-coefficient (Tc) based similarities between the DPCs and approved drugs listed in DrugBank, constructed at 40% similarity cut-off on Tc value. An edge (represented as green line) connects a DPC-drug pair if the Tc value between them is ≥0.4. Of the total 30 DPCs of *T. purpurea*, 27 DPCs (as red triangular nodes) are found to pass the similarity criterion with 75 drugs (purple octagonal nodes) *via* 211 PC-DR pairs. The edge width between 211 pairs in the network varies according to their Tc value, where thicker width represents high value of Tc and thinner represents lower value of Tc. The size of the nodes varies according to their degree centrality value in this network.

TPU_26 (Fraxinol) which shows similarity with an approved drug methoxsalen has been suggested as an uvioresistent agent for its application in UV cosmetics when considered in its glucoside form (CN104367486A, 2015). This shows that the plant *T. purpurea* is a significant source for providing active compounds that could be of considerable importance in future drug-design approaches.

### 3.4 Phytochemical-protein target (PC-PT) network

To ascertain the polypharmacological action of PCs present in *T. purpurea*, interactions between PCs against their 542 PTs are represented in the form of an interaction network (**referred as PC-PT network**) **(Figure 4(A); Supplementary Table-6)**.For one phytochemical corresponding to TPU_52 (Apollinine), no protein targets could be screened-in against the selection criteria adopted for protein-target identification. Therefore, the network corresponds to interactions between 75 PCs and their 542 PTs with network-size of 617 nodes (30DPCs + 45PCs + 542 PTs) and 1,769 edges. The network is analyzed for its degree centrality measure (*C*_*d*_) which is defined as the number of other nodes to which a particular node is connected.

**Fig. 4.**
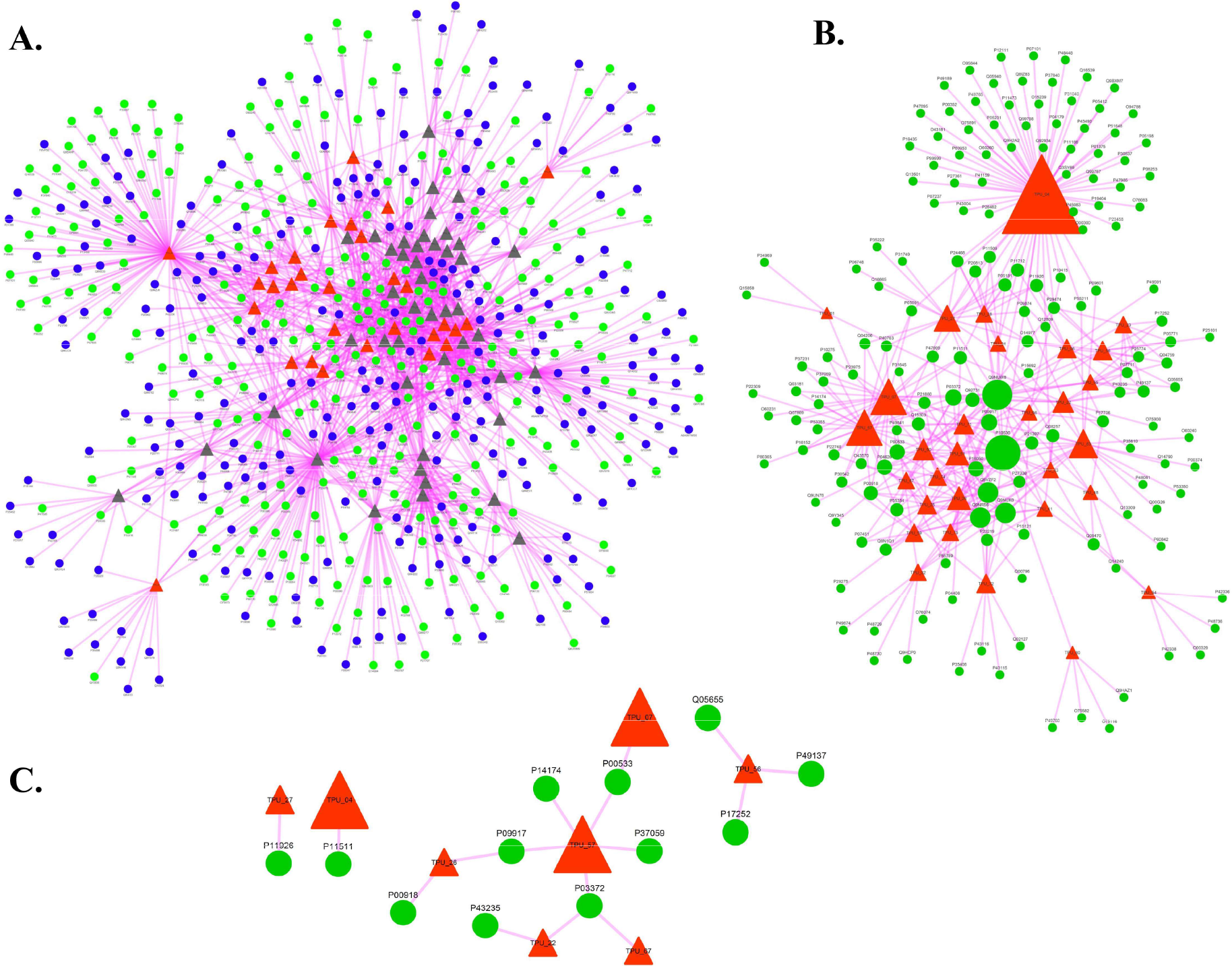
**A: Phytochemical-protein target (PC-PT) network;** PC-PT network shows the association of 75 phytochemicals (PCs, represented as triangular nodes) of *T. purpurea* with their 542 protein targets (PTs, represented as circular nodes). The PC-PT network consists of 617 nodes and 1,769 edges. For the ease of differentiation, phytochemicals (represented as grey coloured nodes) corresponding to the class of DPCs (*i.e.* 30 DPCs) are coloured differently (as red triangular nodes). Here, 161 PCOS proteins are highlighted in the network using green coloured circular nodes. Among all the PCs, TPU_04 holds the maximum connection with the protein targets *i.e*. 105. For the phytochemical having id TPU_52, no protein targets could be identified therefore the networks consist of interactions of only 75 PCS and their 542 protein targets. **B: DPC-PP network**; DPC_PP network represents the sub-network of PC-PT network corresponding to the interactions between 30 DPCs (red triangular nodes) and 161 PCOS proteins (green circular nodes). The size of the nodes varies according to their degree centrality value in the DPC-PP network, where largest node size of connection of TPU_4 shows that among other DPCs it possess the highest regulatory potential by targeting 68 PCOS proteins. **C: High-confidence interaction (HCI) network;** High confidence interactions among the DPCs and PCOS proteins were derived from DPC-PP network and a sub-network is constructed that is referred as HCI network. The HCI interaction were chosen on the basis of the interactions predicted by at-least two of the three protein-target prediction algorithms used in this study.

The (*C*_*d*_) values of 105 and 106 corresponding to TPU_04 and TPU_05, respectively, highlight the high regulatory potential of these PCs compared to others in the list. Among the list of drugable phytochemicals, TPU_04 ranks as the top PC due to its high polypharmacological actions. As already discussed about the investigations on TPU_04 (rotenone) as an experimental drug, the high polypharmacological activity of the same further adds up to its importance.

To understand the phytochemical level effects of *T. purpurea* against PCOS, the interactions between PCs and PCOS proteins were further focused. For this, a sub-network specific to the DPCs of *T. purpurea* and proteins corresponding to PCOS gene library was derived from PC-PT network. The obtained network consisted of 400 interactions among 161 PCOS-proteins and 30 DPCs (**DPC-PP Network; Supplementary Table-7; Figure 4(B)**). When analyzed for another important network centrality measure known as betweenness centrality (*C*_*b*_), TPU_04 returned the maximum value of 0.503 as compared to other DPCs. The high value signifies the importance of TPU_04 as a communicating agent in the overall network, highlighting its global information spreading capacity (Girvan & Newman, 2002).

Other DPCs for which the *C*_*b*_ value is high and therefore demand attention for detailed investigations are TPU_63, TPU_07 and TPU_57 with *C*_*b*_ value 0.176, 0.096 and 0.093, respectively. When checked in the PC-DR network, to examine their existing similarity with DrugBank drugs, all of these DPCs (TPU_04, TPU_63, TPU_07 and TPU_57) were successfully mapped. This further strengthens the lead like (drug-like) properties of these DPCs for their future implication in the disease management.

High-confidence interaction pairs (HCI pairs) defined as pairs predicted by at-least two of the three considered protein-target prediction algorithms were also checked for their presence in the network. This is essential to identify high confidence regulatory DPCs so that they can be put forward for their detailed interaction studies with their respective protein targets and other *in-vitro* studies. In the DPC-PP network, 17 HCI pairs corresponding to the interactions of 8DPCs with 12 PCOS proteins are identified and highlighted in the network (**Figure 4(C); Supplementary Table-7**). It is interesting to note that among the above mentioned 4 DPCs highlighted at the level of betweenness centrality evaluation, 3 (TPU_04, TPU_57 and TPU_07) are constituent of the HCI pairs, also. We believe that these DPCs could be the key regulators of the PCOS proteins for which detailed investigation could be followed up to screen novel regulatory DPCs for PCOS.

### 3.4 Phytochemical, protein targets and biological pathways network (PC-PT-BP)

Drugs shown to target multiple proteins have the ability to regulate more than one disease-associated pathway. Thus, to explore the multi-pathway regulating effects of the *T. purpurea*, all the protein targets of PCs were mapped to the KEGG database. The information of the identified pathways was incorporated into the PC-PT network data to construct a tripartite network of PC-PT-BP consisting of 624 nodes (76 phytochemicals, 542 protein targets, 6 pathways) and 2,759 edges **(Figure 5A**). 270 unique human pathways, distributed among 6 categories *i.e.* metabolism, genetic information processing, cellular processes, organismal systems and human diseases, were shown to be regulated by the PCs of *T. purpurea*. The detailed mapping of all the PTs into different human pathways is given in **Supplementary Table-8**.

**Fig. 5.**
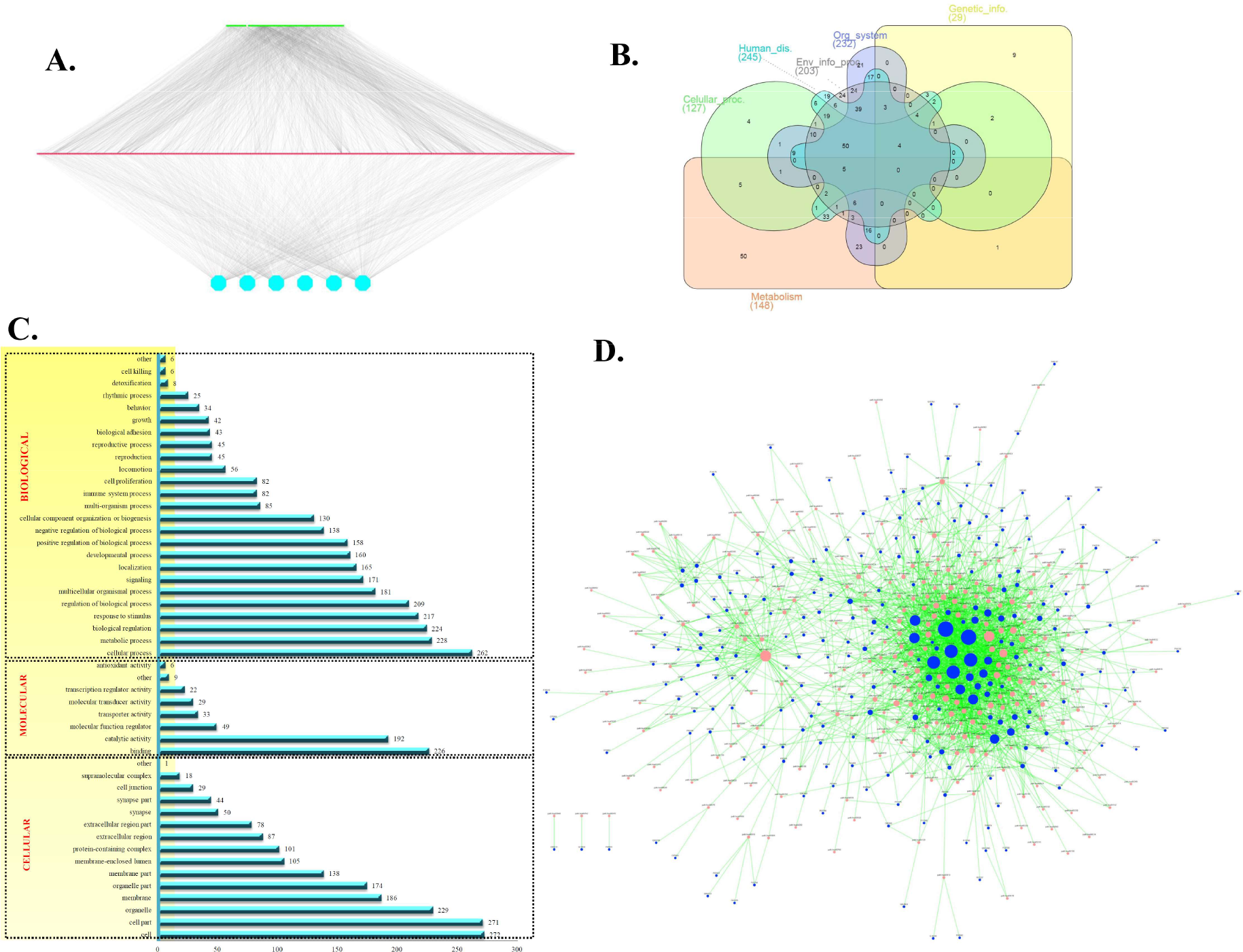
**A: Phytochemical, protein-target and biological pathway (PC-PT-BP) network;** A tripartite network of PC-PT-BP consisting of 624 nodes (76 phytochemicals, 542 protein targets, 6 pathways) and 2759 edges. The top green layer represents 76 phytochemicals and middle red layer represents their 542 protein targets. Protein targets are grouped into six KEGG-pathway categories *viz*, metabolism, genetic information processing, cellular processes, organismal systems and human diseases; each group is represented as a blue colored node at the bottom layer of the network. **B: Distribution of the protein-targets of *T. purpurea* among KEGG pathway classes;** The distribution of 542 PTs of *T. purpurea* among six KEGG-pathway classes *i.e.* metabolism, genetic information processing, cellular processes, organismal systems and human disease, each differentiated using different color. The maximum number of protein targets *i.e.* 245 were associated with the class of “human diseases” while the minimum number of targets *i.e*. 29 with the “genetic information” class. The distribution of proteins is presented in the form of Venn diagram, generated using InteractiVenn (http://www.interactivenn.net/).**C: GO based enrichment analysis of protein targets of DPCs;** Gene-ontology based enrichment of the 278 protein targets of 30 DPCs of *T. purpurea* were found to be distributed among 15 categories of Cellular component, 8 Molecular function and 25 Biological processes. The length of the blue-bar corresponds to the number of protein targets associated with that particular GO-term, where majority of the protein targets are associated with the cell and cellular process, followed by organelle association and metabolic processes with 272, 262, 229 and 228 proteins, respectively. **D: PCOS specific protein target-biological pathway (PT-BP) network;** A sub-network of the PC-PT-BP network, specific to the proteins associated with PCOS (obtained from PCOS-pool) and their associated KEGG pathways specific to *Homo sapiens*. The blue coloured nodes represent the PCOS-proteins and their associated human-pathways are presented using peach coloured nodes, where size of the nodes varies according to their degree value in the network. Metabolic pathway (hsa01100) and cancer-associated pathway (hsa05200) were found to possess the largest node-size amongst all the pathways, representing the maximum number of PCOS-proteins in association (that correspond to 61 PCOS proteins with hsa01100 and 57 with hsa05200).

The PC-PT-BP network when analyzed in detail, shows that multi-pathway effects are exhibited by the PTs of *T. purpurea* where 542 PTs are distributed among the 78 pathways of human disease, 74 of organismal system, 15 of cellular process, 11 of genetic information, 28 of environmental information processing and 59 of metabolism category with 245, 232, 127, 29, 203 and 148 proteins, respectively (**Figure 5B**). The distribution of the proteins presented in the form of Venn diagram was generated using InteractiVenn (http://www.interactivenn.net/) (Heberle, Meirelles, da Silva, Telles, & Minghim, 2015). Among the class of human diseases, majority of the proteins were involved in cancer (hsa05200; pathways in cancer). Similarly, in organismal system pathway, osteoclast differentiation and chemokine signaling (hsa04380 and hsa04062 respectively); apoptosis (hsa04210) under cellular processes; PI3K-Akt signaling pathway (hsa04151) under environment information processing; and protein-processing in endoplasmic reticulum (hsa04141) under genetic information processing class were highly enriched. The results are in line with the findings where involvement of cancer (Dumesic & Lobo, 2013), osteoclast differentiation (Krishnan & Muthusami, 2017), apoptosis (Yan et al., 2012) and PI3k-Akt signaling pathway (Li et al., 2017) are found to be in close relationship with PCOS. The anti-PCOS effects of *T. purpurea* can be directly linked to these proteins through the protein targeting capability of its DPCs.

To elucidate the cellular, molecular and biological level effects of *T. purpurea*, protein targets of this plant were subjected to gene ontology enrichment analysis. 278 protein targets of 30 DPCs were found to be associated with 3,762 unique GO terms. The protein targets of *T. purpurea* were found to be distributed among 15 categories of Cellular component, 8 categories of Molecular function and 25 categories of Biological process **(Figure 5C)**. Majority of the protein targets were associated with cell and cellular process, followed by organelle association and metabolic processes. The GO enrichment analysis revealed that *T. purpurea* possesses the ability to control the diverse functions of a human cell through their protein targets.

To spot the pathways targeted by PCOS proteins, a network specific to PCOS proteins and their regulatory human pathways referred to as **PCOS PT-BP network** was derived and analyzed for the enrichment analysis (**Figure 5D**). It was interesting to note that the highly enriched pathways in the network (**Figure 5D**) corresponds to the metabolic pathway (hsa01100), cancer-associated pathway (hsa05200) and PI3-Akt signaling (hsa04151) owing to 61, 57 and 40 PCOS proteins, respectively. The role of metabolic pathway and cancer-associated pathways in PCOS is well studied as the prevalence of metabolic syndrome accounts for 37.5% (Mandrelle et al., 2012) while the development of endometrial cancer is 2.7 fold in women with PCOS (Dumesic & Lobo, 2013).

Although the association of PI3-Akt signaling in PCOS is well-studied from past several years, the effectiveness of traditional medicinal plants and plant-based formulations for their PI3K-AKT targeting capacity is still unexplored. Towards the exploration of beneficial effects of *T. purpurea* in PCOS through PI3k-AKT signaling pathway, the protein targets of the plant (PTs of DPCs) were mapped to this pathway. As seen in the **Supplementary Figure 1**, it was observed that the proteins got mapped at **20** locations in this pathway, which suggests that the phytochemicals of this plant may exert substantial control over the pathway.

### 3.5 Disease-association network construction and analysis

To infer about the regulatory effects of the *T. purpurea* against various human diseases, the protein targets of its constituent phytochemicals were searched for their disease association. Out of 542 PTs of 76 PCs, 446 PTs were found to be associated with at least one disease. It was interesting to note that the PTs of *T. purpurea* were involved in 2,918 different diseases, broadly classified into 25 categories, based on MeSH disease class (**Supplementary Figure 2**). The highest numbers of protein targets were shown to be associated with Neoplasm and Nervous system diseases with 281 and 236 proteins, respectively. The anti-cancer properties of this plant on MCF-7 cell lines have demonstrated its efficacy in cancer management (Gulecha & Sivakuma, 2011). The high association of the P01375 (TNF; Tumor Necrosis Factor) with 343 diseases reflect the relative importance of this protein as a potential target. Compared to normal patients, the high concentration of TNF-α has been linked to observed androgen access and insulin resistance in PCOS patients (Gao, Gu, & Yin, 2016). When checked for the regulatory DPCs against P01375, the association of the proteins with TPU_04 (Rotenone) came into light.

Since we are also interested in exploring the effectiveness of *T. purpurea* against the PCOS comorbid diseases, proteins constituting the PCOS-pool alongwith the derived disease associations were used to develop a **PP-DA**(PCOS-protein—Disease-association) network (**Figure 6**).

**Fig. 6:**
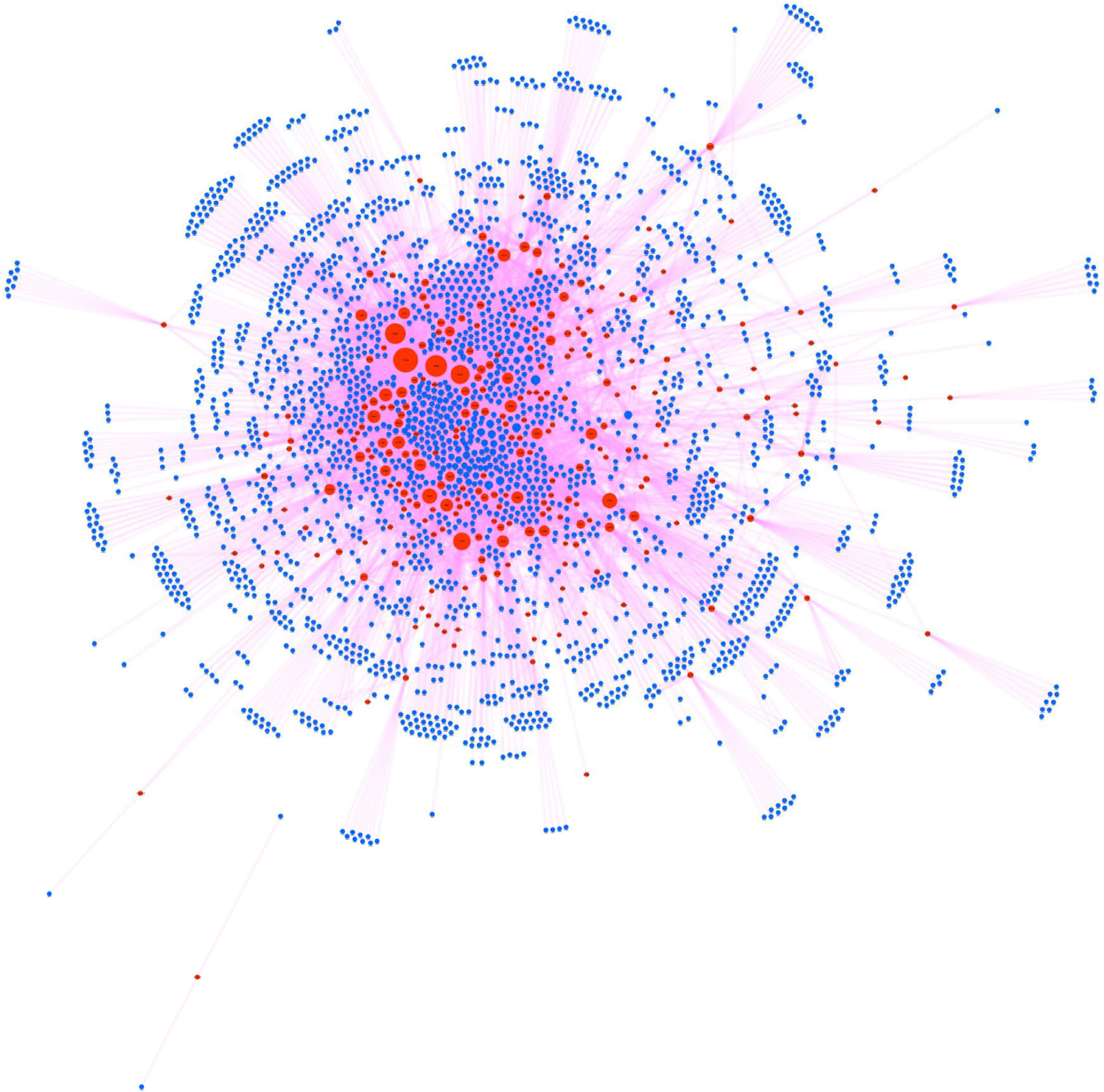
PP-DA (PCOS-protein—Disease-association) network; The network showing the disease association of 261 PCOS proteins (red coloured nodes) with 2,403 diseases mentioned in MeSH (blue-coloured nodes). Network consists of 2,664 nodes (261 PCOS proteins and 2,403 disease classes) and 9,343 edges, where schizophrenia (a mental disorder, C0036341) is found to share the maximum number of PCOS-proteins *i.e.* 343. The size of PCOS proteins in the network varies according to their connections with the disease class *i.e.* degree value.

As seen in PP-DA network having network size of 2,664 nodes (261 PCOS proteins and 2,403 disease classes) and 9,343 edges, schizophrenia (a mental disorder, C0036341) is found to share the maximum number of PCOS-proteins *i.e*. 343. Although the direct association of this disease with PCOS is less explored, a large number of studies have related PCOS with various mental as well as psychiatric disorders (Blay, Aguiar, & Passos, 2016; Hung et al., 2014). While the PCOS genes are associated with 2,403 diseases, high degree of interconnections apart from the schizophrenia were found with C0006142 (breast cancer), C0376358 (prostate cancer), C0033578 (prostate cancer) and C3714756 (Intellectual disability/ mental retardation) *via* 69, 63, 63 and 56 PCOS proteins, respectively. Detailed mapping of the PCOS proteins into various disease classes is given in the **Supplementary Table 9**.

Our specific investigations in the form of two case studies are detailed in the following subsections.

#### Case study-I: Investigation of regulatory DPCs effective against PCOS comorbidities

Literature data suggests that the association of PCOS with neurological disorders, especially; bipolar disorder (BPD), depression and anxiety are of high confidence. Earlier reported studies provided the evidence that the risk of depression among the women affecting with PCOS is linked to insulin resistance, where an elevated level of insulin has been observed in the patients when assessed by HOMA-IR (Homeostasis Model Assessment of Insulin Resistance) (Greenwood, Pasch, Shinkai, Cedars, & Huddleston, 2015). Also, a cross-sectional based observation study shows that the severity of symptoms associated with anxiety is much more prevalent in PCOS patients, compared to depression (Deeks, Gibson-Helm, & Teede, 2010). Comorbid association between PCOS and psychiatric disorders like BPD (Bipolar disorders) has been observed at both pharmacotherapy dependent (especially valproate treatment) (McIntyre, Mancini, McCann, Srinivasan, & Kennedy, 2003) and valproate independent PCOS-BPD association studies (Klipstein & Goldberg, 2006). To screen potential multi-targeting and synergistic regulators from the *T. purpurea* against the protein targets of these diseases, the following approach was adopted.

Firstly, high confidence disease associations of protein targets were screened from DisGeNET using the threshold score of > 0.05. Gene-disease association (GDA) score (*S*) by DisGeNET ranges in between 0-1, in which both data-mining and literature-based associations are being taken into consideration while calculating the score. As described by DisGeNET, the score (*S*) is given by;

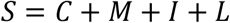

Where, *C* component takes into account of curated data, *M* with animal model data, *I* with inferred data and *L* corresponds to literature data supporting the GDA. The threshold of 0.05 was chosen so as to include all the associations having non-zero score in either C or M or I or at least 5 publications (*i.e*. *L* content of the score), where each publication contributes to the score of 0.01 leading to an overall score of 0.05. This threshold in the score excludes all the GDAs that have no support from C or M or I as well the GDAs having 5 or less number of publications support from the text-mining results. The detailed description of the scoring schema, its components and calculations may be checked at http://www.disgenet.org/dbinfo#score.

Using this criterion, 579, 235 and 740 proteins were selected corresponding to BPD, anxiety and depression, respectively. Using a similar strategy, 223 proteins specific to PCOS were chosen for further analysis. The proteins were mapped into the PC-PT network to identify their potential regulators from the drug-like phytochemical pool of *T. purpurea*. Upon mapping, 30 DPCs were found to interact with 83 proteins associated with the above discussed four classes of diseases using 214 compound-protein pairs (**Figure 7A**). The multi-disease associations of a protein were observed at this point, where multiple proteins overlap between the 4 classes of diseases discussed here as shown in **Figure 7B**.

**Fig. 7:**
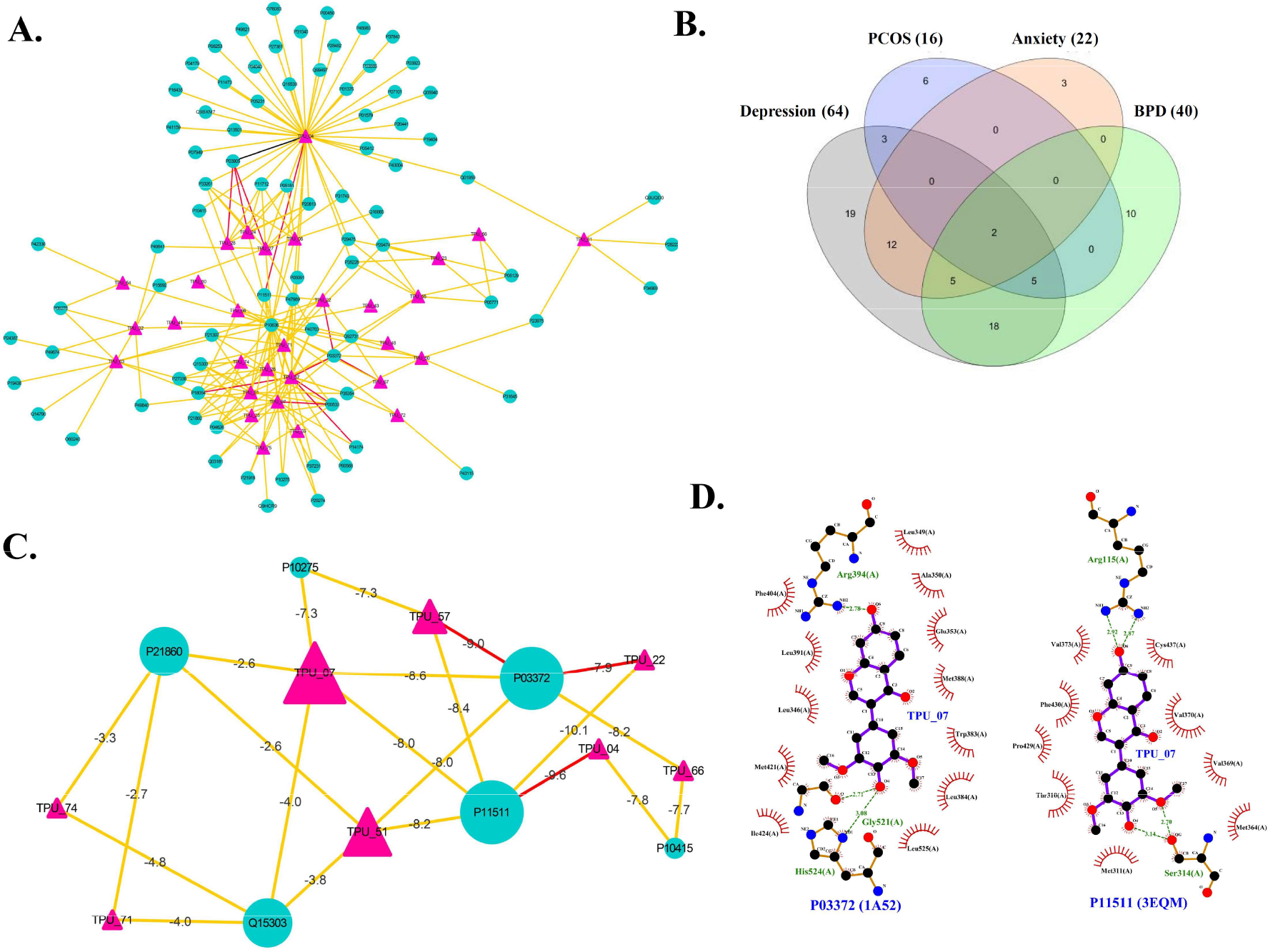
Investigation of regulatory DPCs effective against PCOS comorbidities; **A:** A sub-network of PC-PT network, specific to the proteins associated with PCOS and its three comorbities *i.e*. bipolar disorder (BPD), depression and anxiety. 30 DPCs (pink triangular nodes) can target 83 proteins (green circular nodes) using 214 phytochemical-protein pairs. The phytochemical-protein pairs are represented using differently colored edges, where black color corresponds to the pair predicted by all the 3 target prediction algorithms, red and yellow colored edges correspond to pairs predicted by at-least two and at-least one target prediction algorithms, respectively. **B:** The Venn-diagram showing the distribution of 83 protein targets of 30 DPCs into PCOS, bipolar disorder (BPD), depression and anxiety, where multiple proteins show their overlap amongst the 4 disease classes. **C:** A sub-network specific to 8 DPCs highlighting their multi-targeting and synergistic action against 6 protein targets associated with PCOS-comorbidities, where proteins are represented using green colored circular nodes and their regulatory DPCs with pink triangular nodes. Docking energy values of each interaction pair in the network is presented along the edges; maximum affinity is shown by TPU_22 for P11511 having binding energy value of −10.1 kcal/mol. **D:** Ligplot interaction of the docked complex of TPU_07 with P03372 (Estrogen receptor) and P11511(Aromatase enzyme), where both hydrogen bonds and hydrophobic interactions are involved in stabilizing the protein-ligand complex. Hydrogen bonds are represented using dashed lines while the residues involved in hydrophobic interactions as arcs.

The proteins specific to each disease class as well as those overlapping are given in (**Supplementary Table 10**). For the identification of DPCs that may act on multiple scales, the protein targets of DPCs were mapped in the **PP-DA** network and proteins with the multi-disease association, considering the above 3 comorbid diseases were taken for further analysis. In this manner, 7 classes of proteins comprising 10 proteins were obtained and the number of proteins belonging to each class is given in **Table 2**.

**Table 2:** Distribution of the proteins among different classes of comorbid diseases

| Serial No. | Class | No. of proteins | Proteins |
| --- | --- | --- | --- |
| 1. | PCOS and Depression | 3 | P10275, P41159, P11511 |
| 2. | PCOS and Anxiety | 0 | ---- |
| 3. | PCOS and BPD | 0 | ---- |
| 4. | PCOS, Depression and Anxiety | 0 | ---- |
| 5. | PCOS, Anxiety and BPD | 0 | ---- |
| 6. | PCOS, Depression and BPD | 5 | P10415, P21860, P05231, P03372, P07101 |
| 7. | Associated with all four (PCOS, Depression, Anxiety and BPD) | 2 | P01375, Q15303 |

The selected 10 proteins were mapped to PC-PT network to identify their regulatory DPCs. The 10 proteins were found to interact with 12 DPCs *via* 30 interactions. The network was analyzed in detail to identify multi-targeting and synergistic regulators. The investigations led to the screening of 8 DPCs against 6 protein targets (**Figure 7C**). Each interaction of the network is subjected to molecular docking analysis to identify their potential binding energy values **(Table 3)**.

**Table 3:** Docking analysis of the phytochemical-protein targets pairs specific to multi-targeting and synergistic regulation.

| S.NO. | PC_ID | Protein target |  | Binding energy (Kcal/mol) |
| --- | --- | --- | --- | --- |
|  |  | Uniprot ID | PDB ID |  |
| 1. | TPU_04 | P11511 | 3EQM | -9.6 |
| 2. | TPU_04 | P10415 | 1E3G | -7.8 |
| 3. | TPU_07 | P03372 | 1A52 | -8.6 |
| 4. | TPU_07 | P11511 | 3EQM | -8 |
| 5. | TPU_07 | P10275 | 1E3G | -7.3 |
| 6. | TPU_07 | Q15303 | 3BCE | -4 |
| 7. | TPU_07 | P21860 | 1M6B | -2.6 |
| 8. | TPU_22 | P11511 | 3EQM | -10.1 |
| 9. | TPU_22 | P03372 | 1A52 | -7.9 |
| 10. | TPU_51 | P11511 | 3EQM | -8.2 |
| 11. | TPU_51 | P03372 | 1A52 | -8 |
| 12. | TPU_51 | Q15303 | 3BCE | -3.8 |
| 13. | TPU_51 | P21860 | 1M6B | -2.6 |
| 14. | TPU_57 | P03372 | 1A52 | -9 |
| 15. | TPU_57 | P11511 | 3EQM | -8.4 |
| 16. | TPU_57 | P10275 | 1E3G | -7.3 |
| 17. | TPU_66 | P03372 | 1A52 | -8.2 |
| 18. | TPU_66 | P10415 | 1E3G | -7.7 |
| 19. | TPU_71 | Q15303 | 3BCE | -4 |
| 20. | TPU_71 | P21860 | 1M6B | -2.7 |
| 21. | TPU_74 | Q15303 | 3BCE | -4.8 |
| 22. | TPU_74 | P21860 | 1M6B | -3.3 |

As seen in **Table 3**, it was observed that the maximum binding energy value of −10.1 kcal/mol exists between P11511 and TPU_22. The DPC TPU_22 is also capable of regulating the P03372 with a good binding energy value of −7.9 Kcal/mol. The protein P03372 (Estrogen receptor; ESR1gene) is known to be associated with above discussed 3 diseases *i.e.* PCOS, BPD and depression.

Previous studies have correlated the role of ESR1 gene in PCOS *via* estrogen metabolism (Jiao et al., 2018). Depression-like behavior in postpartum rats was shown to be associated with estrogen-mediated metabolism (Furuta et al., 2013). While in BPD subjects, ESR1 is shown to be associated with behavioral traits (Pinsonneault et al., 2017). This suggests that the role of TPU_22 is highly noteworthy in dealing with the co-morbidities associated with PCOS.

When checked in the phytochemical-drugs similarity network (PC-DR), the compound shows its similarity with 12 drugs, which further strengthens the drug-like potential of this phytochemical. Along with TPU_22, TPU_51 and TPU_57, TPU_07, TPU_66 on the ESR1 proteins show the binding energies in the range of −7.9 Kcal/mol to −9.0 Kcal/mol. We propose these compounds for future *in-vitro* as well as *in-vitro* studies to decipher the synergistic actions of these molecules to get detailed insights about their mechanism of action to further strengthen the anti-PCOS effects of the *T. purpurea*. Also, the multi-targeting potential of TPU_07 is worthy of attention, where the compound can regulate 5 proteins of the above considered 10 proteins. When checked in detail for the molecular interactions of TPU_04 with its interacting proteins, the phytochemical shows good binding energy value of −8.6 and −8.0 with P03372 and P11511, respectively. As already discussed about the association of P03372 with PCOS, BPD and depression, the protein could be considered as a potential target of TPU_07. Interaction analysis of the docked complex of TPU_07 with P03372 shows that the phytochemical holds the ability to form hydrogen bond with the protein at arginine (Arg 394), glycine (Gly 521) and histidine (His 524) residues.

As seen in **Figure 7D**, the protein-ligand complex is also stabilized by the hydrophobic interactions governed by 12 residues of the protein. When checked for the molecular interactions of TPU_07 with P11511, the formation of strong H-bond governed by arginine (Arg115) and serine (Ser314) residues further strengthens the molecular interaction between the protein-ligand complex, leading to a stable interaction pair with P11511. Binding energy of the protein (P11511) shows that the protein possesses the ability to interact with TPU_07 with 8.0 kcal/mol of energy. The protein (P11511; aromatase enzyme) is associated with PCOS and depression **(as seen in Table-2),** where the usage of aromatase inhibitors are taken into consideration while dealing with the complications associated with PCOS. Although, not opted as first-line therapy, the aromatase inhibitors are good choice of ovulation-inducing agent (OIA) for their lowered side effects compared to the well-known oral OIA, clomiphene citrate (Misso et al., 2012). This suggests that TPU_07 could be of promising role as a potential aromatase inhibitor and the investigation require experimental support from *in-vitro* and *in-vivo* inhibition studies.

In the similar manner, the network can also be checked for DPCs that may work in synergism to regulate a protein of interest. The involvement of the protein targets of *T. purpurea* in diverse class of diseases shows that the plant possesses the ability to work on multiple scales, highlighting the ethnopharmacological importance of the plant.

#### Case study II: Construction of PCOS specific PPI network (PCOS-PIN) and examining its modules regulated by *T. purpurea*

To ascertain the comprehensive interactions of PCOS genes on the entire human system, a PPI network specific to PCOS proteins was constructed and analyzed in detail. The construction of PCOS-PIN follows the development of a high-confidence human-PPI from STRING database with the confidence score of ≥ 0.9. A total of 7,972 PCOS proteins were searched in the human-PPI, resulting in the mapping of 4,456 proteins. The 4,456 proteins were having the tendency to interact with each other using 35,197 interactions, the network corresponding to these interactions constitute **PCOS-PIN (Figure 8A, Supplementary Table 11)**. The network analysis of the PCOS-PIN shows that node-degree distribution follows a power-law distribution with *y* = 3014.9*x*^−1.520^ (**Figure 8B**).

**Fig. 8:**
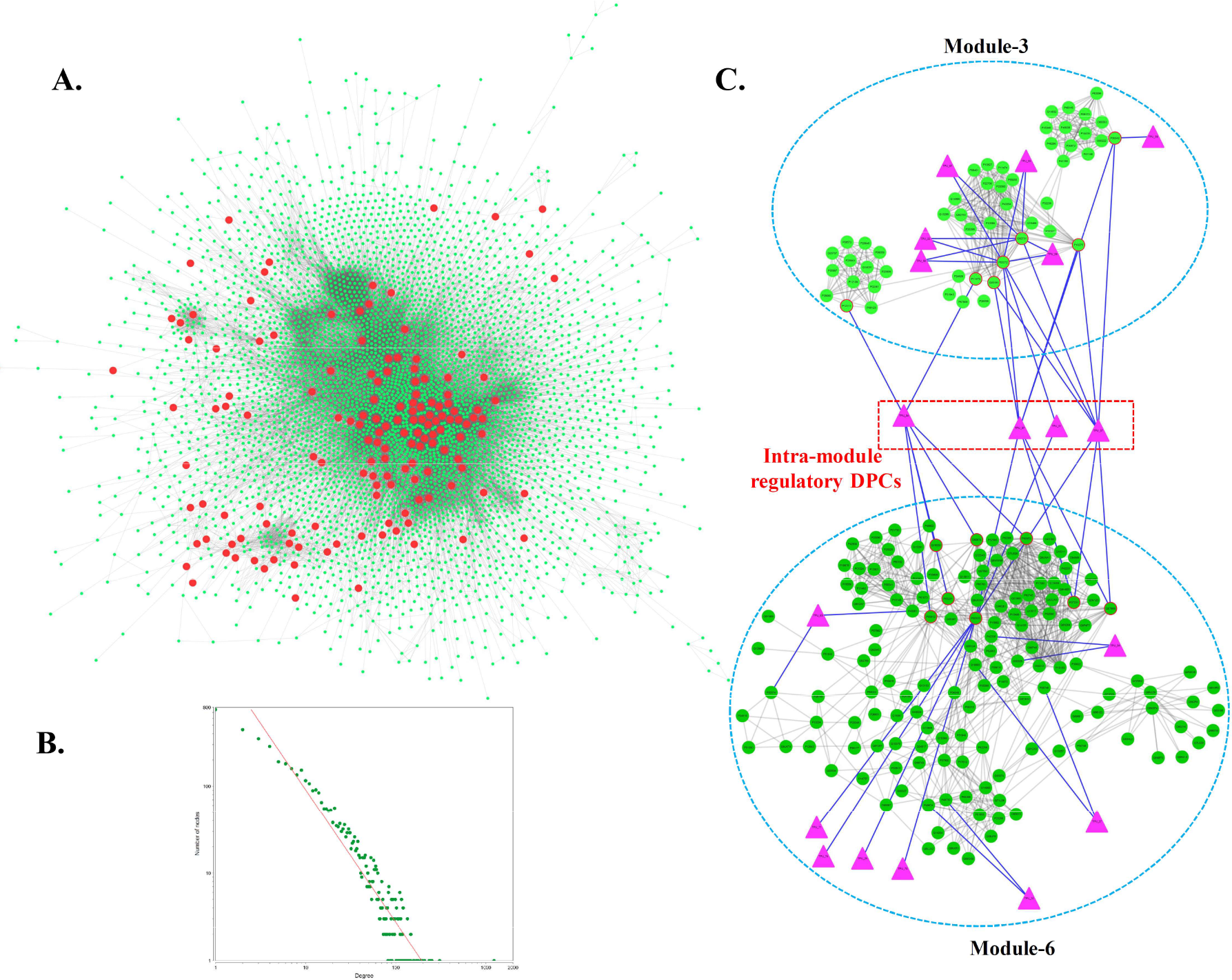
Regulatory potential of *T. purpurea* on PCOS-PIN and its constituent modules; **A:** High-confidence sub-network of human PPI generated using the STRING score ≥ 900, specific to PCOS proteins. The network consists of 4,456 proteins and 35,197 edges. The red colored circular nodes represent the location of protein targets of *T. perpurea* in the PCOS-PIN. **B:** Node-Degree distribution of the PCOS-PIN. X and Y-axes of the graph represent the number of nodes and their respective degree values in the PCOS-PIN, where both the axes are plotted on the logarithmic scale. **C:** Multi-module regulatory potential of the phytochemicals of *T. purpurea* against two modules (Module-3 and Module-6) are shown with the constituting protein targets (green colored circular nodes) and their regulatory DPCs (pink colored triangular nodes). Four intra-module regulatory DPCs (TPU_04, TPU_07, TPU_51 and TPU_57) are highlighted in this figure in red colored triangular box, where these 4 DPCs possess the tendency to directly target 7 and 8 proteins (shown in red outlined circles) of module-3 and module-6, respectively.

Since proteins tend to work together in a group, therefore the obtained network was subjected for the cluster (module) analysis to identify potential clusters and their associated proteins using the MCODE algorithm (Bader & Hogue, 2003). Module analysis of the network shows that PCOS-PIN is regulated by multiple modules, where proteins associated with each module or cluster work in a coordinated manner to provide structural and functional integrity to the PIN. Top 10 modules with dense connections were taken for the subsequent analysis and were subjected for GO-based annotation using DAVID (Dennis et al., 2003). This is essential to identify the functional role of the clusters in the human system in relation to PCOS. This also aids towards understanding the complex inter-relationship between the genes associated with a particular cluster and their functional relevance in the pathogenesis of PCOS.

Functional association of the clusters shows that proteins of the PCOS-PIN network are involved in the regulation of various cellular events, the majority being associated with signaling events of the cell or signal transduction pathways *i.e*. module-2, 3, 5, 6 and 7. As given in **Supplementary Table-12**, it can also be observed that PCOS proteins also exert their effects on regulating apoptotic process *via* module-8. Altered expression of genes associated with apoptotic process has been observed in the endometrium of the women suffering from PCOS, a study from China reports that the level of bax-gene is significantly higher in PCOS compared to normal levels (Maliqueo et al., 2003; Wu et al., 2017). The participation of the PCOS protein in the regulation of other cellular events like rRNA processing, mitochondrial electron transport, response to DNA damage and transcription regulation could be linked to the proteins of module 1, 4, 9 and 10, respectively.

As the case-study aims towards examining the module regulatory potential of the phytochemicals of *T. purpurea*, the proteins associated with a protein-target list of 30 DPCs (*i.e.* 278 proteins) were searched for their participation in the modules. Mapping of 278 proteins onto the considered 10 highly clustered modules could help us to identify the high regulatory role of the DPCs towards module-3 and module-6 as these modules are rich in protein associated with the protein-target list of DPCs. Majority of the proteins of module-3 are associated with “steroid hormone-mediated signaling pathway” where irregular steroid hormones level (Doi, Al-Zaid, Towers, Scott, & Al-Shoumer, 2005), altered steroid metabolism (Oróstica et al., 2016) and over expression of steroid receptors (Maliqueo et al., 2003) have been observed in the cases of PCOS. Proteins of module-6 show their participation in the “signal transduction” event. The role of association of various signaling events like PI3k-Akt (Li et al., 2017; Makker, Goel, Das, & Agarwal, 2012), cAMP, WNT, LH and androgen signaling (Dumesic & Richards, 2013) with the pathogenesis of PCOS are well known.

For the detailed investigation about the information of DPCs of *T. purpurea* towards regulating module-3 and module-6, the proteins of both the modules were back mapped in PC-PT network and a sub-network corresponding to module proteins and their regulatory DPCs were constructed. As seen in **Figure 8C**, a total of 18 DPCs have the ability to regulate both the modules, where 10 DPCs can target the proteins of module-3 and 12 DPCs can target module-6. The role of 4 DPCs (TPU_57, TPU_51, TPU_07 and TPU_04) in regulation of both the modules is also highlighted in this figure, where these 4 DPCs possess the tendency to directly target 7 and 8 proteins (**shown in red outlined circles in Figure 8C**) of module-3 and module-6, respectively. Among the 4 DPCs, the phytochemicals corresponding to TPU_04, TPU_07 and TPU_57 should be given special attention, because in addition to their multi-module regulatory effect, the phytochemicals possess high betweenness centrality values in the PC-PT network. The structural similarity of these phytochemicals with the existing drugs of DrugBank, as seen in PC-DR network further supports the need for their detailed investigation towards the design of therapeutically active compounds against PCOS.

In a similar manner, other modules can be studied and checked for the DPCs for their multi-module participation.

#### Summary

Polycystic ovarian syndrome (PCOS) comprises a spectrum of reproductive and non-reproductive complications with a high rate of prevalence amongst the women of reproductive age. Being the major cause of infertility, this syndrome has always gathered the attention of research community for the design of safe and effective treatment measures. The traditional Indian system of healing originated in the Vedic culture of India, holds the information of various herbs towards their effectiveness against PCOS. *T. purpurea,* a well-known Ayurvedic herb is an important component of many Ayurevdic herbal formulations. Various literature studies support the pharmacological importance of the plant towards disease management in humans. However, the multi-targeting pharmacology of its various phytochemicals and their role in dealing with disease-comorbidity is yet to be explored. Hence in this study, the network pharmacology approach of drug discovery has been employed to explore the multi-functional role of phytochemicals present in *T. purpurea* towards managing complex disease comorbities associated with PCOS. The approach works in a hierarchical manner where first step involves the listing of phytochemicals present in *T. purpurea* based on KDD (Knowledge based data discovery) and literature survey method. A list of collected 76 phytochemicals and their chemical information forms the basis of present study. Pharmacological action of *T. purpurea* was studied by predicting the protein-targeting ability of its 76 phytochemicals against 542 human proteins. Only 30 of collected 76 phytochemicals possess the lead-like properties and are termed as drugable phytochemicals (*i.e.* DPCs), identified on the basis of drugability analysis. The therapeutic effects of DPCs in management of PCOS were assessed on the basis of their PCOS-protein targeting capability. For this, a protein-library corresponding to PCOS proteins comprising of 8,199 genes was constructed. The regulatory prospects of 30 DPCs of *T. purpurea* against 161 PCOS-proteins were identified where multi-targeting and synergistic potential of TPU_04, TPU_57 and TPU_07 were prominently highlighted on the basis of degree centrality measure. The multi-targeting aspect of DPCs was examined on the basis of involvement of their protein targets into various human pathways and disease classes where the complex inter-relationship is represented and studied in the form of networks. Detailed analysis of network properties accentuated the high association of metabolic and cancer associated pathways with the PCOS-targeting capability of this herb *. T. purpurea* mediated PI3k-Akt regulation through its 20 protein targets is also examined and presented using pathway mapping studies. The PI3k-Akt pathway acts as a major source of the signaling events towards extracellular response where activation of the same can be correlated with endometrial cancer observed in case of PCOS. Investigation of regulatory DPCs effective to deal the major comorbidities associated with PCOS *i.e.* anxiety, depression and BPD (bipolar disorders) is also highlighted in this work and is discussed as a special case study. 8 potential regulators of *T. purpurea* against 6 PCOS proteins associated with its comorbid diseases were also identified in this work and studied for their docking and interaction analysis. The phytochemicals showing good binding affinity towards their protein targets were also evaluated for their similarity against currently available approved drugs present in DrugBank. TPU_22, TPU_51, TPU_57, TPU_07 and TPU_66 are the compounds being reported in this study to have substantial therapeutic relevance against the PCOS and its associated comorbidities. Augmenting the results obtained in this study with *in-vivo* and *in-vitro* will be of substantial help towards understanding the detailed mechanism of PCOS pathogenesis and design of future PCOS targeting drugs, effective to manage women-health. We believe that network-pharmacology based computational framework developed in this study will be a guiding source towards strengthening the anti-PCOS effect of *T. purpurea* and will also act as a platform in providing a modern scientific outlook to the traditional knowledge of Ayurveda and other traditional medicinal systems.

## Acknowledgements

N.C is grateful to the Indian Council of Medical research (ICMR) for support provided through ICMR-SRF. All the authors thank the support of the Central University of Himachal Pradesh for providing computational facilities.

## Authors’ contribution

V.S. conceptualized and supervised the study. S.C. contributed to data-collection and N.C. performed the data-integration and computational analysis. N.C. and V.S. contributed in the investigations and analysis of results. All the authors contributed in writing of manuscript.

