## Supplementary figures and images for "Deciphering the multi-scale mechanisms of *Tephrosia purpurea* against polycystic ovarian syndrome (PCOS) and its major psychiatric comorbidities: studies from network-pharmacological perspective"

### Supplementary_figure-1.pdf

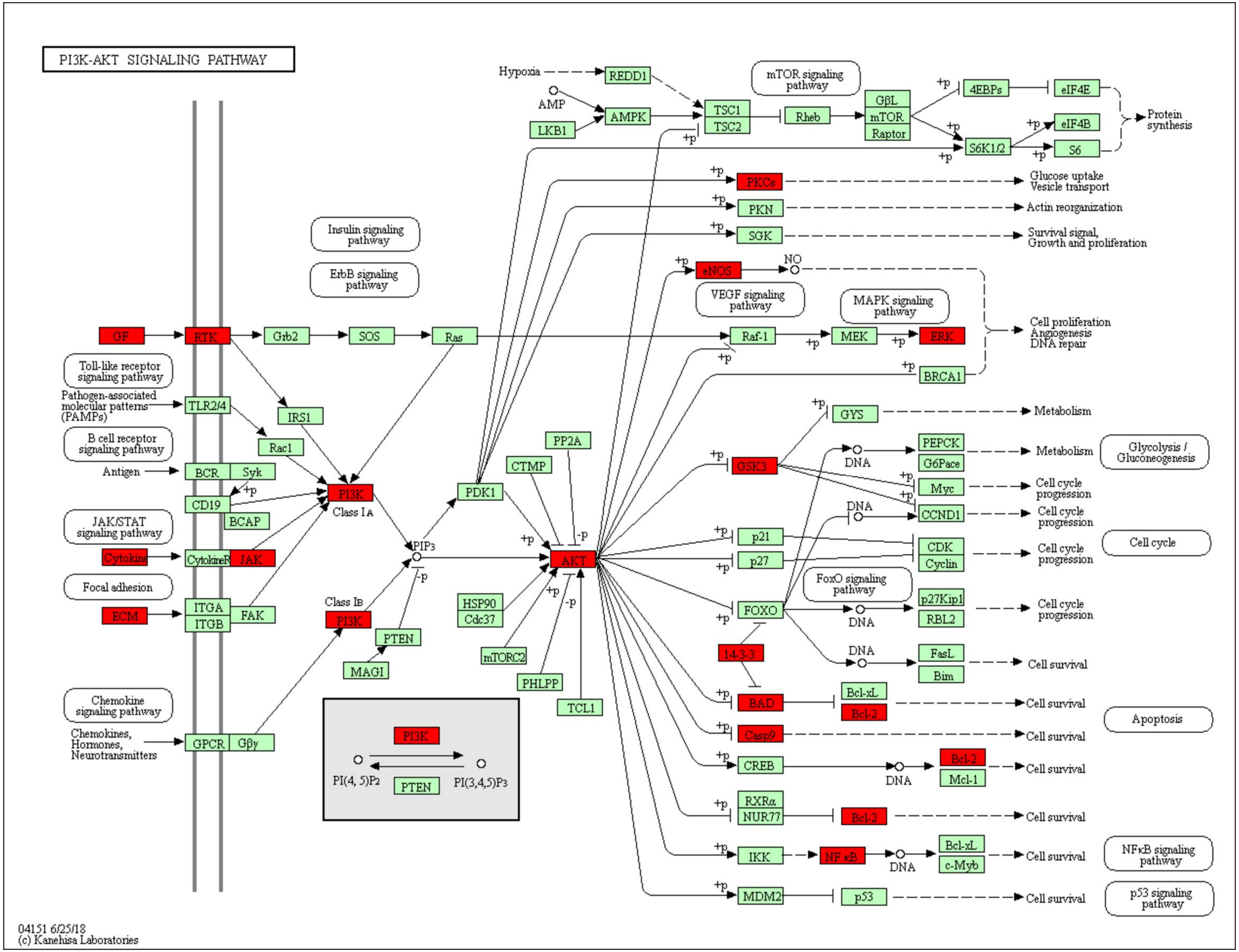

### Supplementary_figure-2.pdf

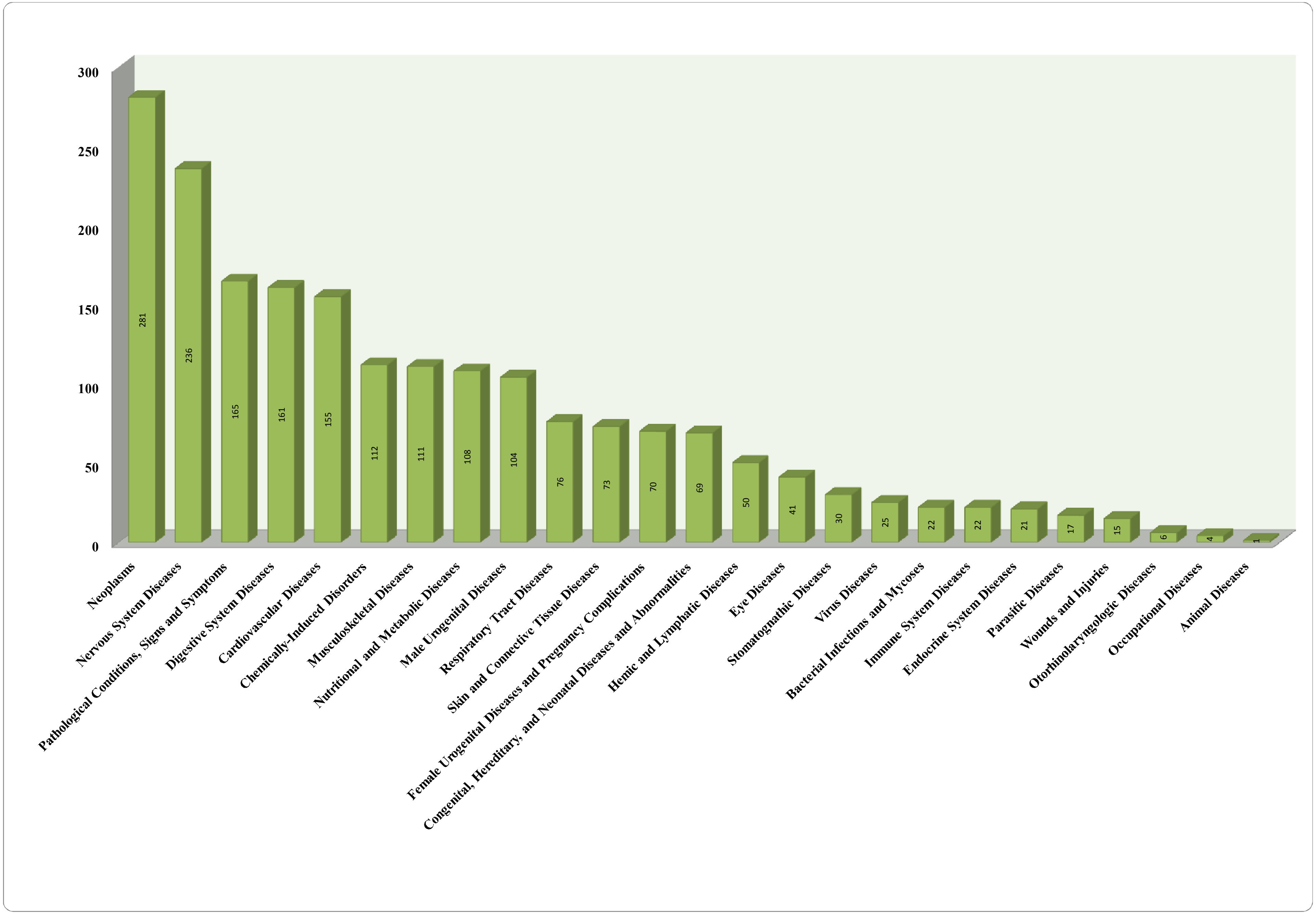
